# Multimodal multilayer network centrality relates to executive functioning

**DOI:** 10.1101/2021.06.28.450180

**Authors:** Lucas C. Breedt, Fernando A. N. Santos, Arjan Hillebrand, Liesbeth Reneman, Anne-Fleur van Rootselaar, Menno M. Schoonheim, Cornelis J. Stam, Anouk Ticheler, Betty M. Tijms, Dick J. Veltman, Chris Vriend, Margot J. Wagenmakers, Guido A. van Wingen, Jeroen J. G. Geurts, Anouk Schrantee, Linda Douw

## Abstract

Executive functioning is a higher-order cognitive process that is thought to depend on a brain network organization facilitating network integration across specialized subnetworks. The frontoparietal network (FPN), a subnetwork that has diverse connections to other brain modules, seems pivotal to this integration, and a more central role of regions in the FPN has been related to better executive functioning. Brain networks can be constructed using different modalities: diffusion MRI (dMRI) can be used to reconstruct structural networks, while resting-state fMRI (rsfMRI) and magnetoencephalography (MEG) yield functional networks. These networks are often studied in a unimodal way, which cannot capture potential complementary or synergistic modal information. The multilayer framework is a relatively new approach that allows for the integration of different modalities into one ‘network of networks’. It has already yielded promising results in the field of neuroscience, having been related to e.g. cognitive dysfunction in Alzheimer’s disease. Multilayer analyses thus have the potential to help us better understand the relation between brain network organization and executive functioning. Here, we hypothesized a positive association between centrality of the FPN and executive functioning, and we expected that multimodal multilayer centrality would supersede unilayer centrality in explaining executive functioning. We used dMRI, rsfMRI, MEG, and neuropsychological data obtained from 33 healthy adults (age range 22-70 years) to construct eight modality-specific unilayer networks (dMRI, fMRI, and six MEG frequency bands), as well as a multilayer network comprising all unilayer networks. Interlayer links in the multilayer network were present only between a node’s counterpart across layers. We then computed and averaged eigenvector centrality of the nodes within the FPN for every uni- and multilayer network and used multiple regression models to examine the relation between uni- or multilayer centrality and executive functioning. We found that higher multilayer FPN centrality, but not unilayer FPN centrality, was related to better executive functioning. To further validate multilayer FPN centrality as a relevant measure, we assessed its relation with age. Network organization has been shown to change across the life span, becoming increasingly efficient up to middle age and regressing to a more segregated topology at higher age. Indeed, the relation between age and multilayer centrality followed an inverted-U shape. These results show the importance of FPN integration for executive functioning as well as the value of a multilayer framework in network analyses of the brain. Multilayer network analysis may particularly advance our understanding of the interplay between different brain network aspects in clinical populations, where network alterations differ across modalities.

**Highlights:**

- Multimodal neuroimaging and neurophysiology data were collected in healthy adults
- Multilayer frontoparietal centrality was positively associated with executive functioning
- Unilayer (unimodal) centralities were not associated with executive functioning
- There was an inverted-U relationship between multilayer centrality and age

## 1. Introduction

A thread of network thinking runs through the history of cognition research. In 1983, Fodor introduced the ‘modularity of mind’ theory of cognition and behavior [1]. He posited that lower-order processes of the mind are modular, with domain-specific modules operating independently without interacting with other modules. Contrastingly, he argued that higher-order cognitive processes such as executive functioning (EF), which is thought to be the most complex and evolutionarily special cognitive domain [2], are global rather than modular. Likewise, the evolution of neuroscience has led to a data-driven approach towards understanding how the brain governs such higher-order cognition by studying the brain as a complex network through the framework of graph theory [3, 4]. Brain regions are thus represented as nodes, and the interactions between them as links.

Different modalities can be used to obtain these brain networks. Anatomically, diffusion magnetic resonance imaging (dMRI) maps the physical connections (i.e. white matter bundles) between the neural elements of the brain, yielding a structural network. Functionally, multiple imaging techniques can be used to observe brain activity. Resting-state functional magnetic resonance imaging (rsfMRI) detects variations in blood oxygenation as an indirect measure of neuronal activity at a high spatial resolution, and magnetoencephalography (MEG) provides a direct measure of the summed electromagnetic activity generated by groups of neurons. In both rsfMRI and MEG, statistical interdependencies between levels of activity in different areas of the brain are used as a measure for functional connectivity [5, 6], yielding functional networks.

The organization of these structural and functional networks appears to be crucial for EF. Although the exact mechanisms underlying this cognitive function remain unknown [7], EF appears to be highly reliant on network integration, i.e. the interplay between specialized modules [8]. Key in facilitating this integration is the frontoparietal network (FPN), a module that plays a crucial central role as a ‘connector’ within the brain network, having diverse connections to other modules of the brain [9]. The network integration that is hypothetically happening in the individual brain regions that form the FPN can be characterized through network measures of centrality. Nodal centrality reflects the relative importance of a node within the network. Highly central regions are typically connected to many other regions, implying a pivotal role in the facilitation of network integration [9, 10]. Indeed, a more central role of the FPN has been related to better EF in unimodal network studies that utilized dMRI [11], rsfMRI [12, 13], or MEG [14].

However, in such unimodal network studies the different aspects of the brain network, e.g. structural and functional, are only studied in isolation, while we know from other types of complex networks that network structure and different types of functional dynamics occurring on top of it jointly and synergistically determine system behavior [15–18]. In the brain, it remains unclear exactly how the integration between these network aspects relates to EF. Nevertheless, the interplay between structural and functional connectivity has been shown to be non-trivial, suggesting both should be considered simultaneously [19–21]. Moreover, in the case of networks based on MEG data, the broadband signal is often filtered into canonical frequency bands, and network analysis is performed for each frequency band separately, but the different imaging modalities and frequency bands each yield unique and even complementary information that should perhaps not be considered in isolation. Unimodal networks are thus limited representations of the essentially multimodal brain network [18, 22, 23]; but until recently we lacked the appropriate tools to integrate multiple modalities into a single network representation.

Multilayer network analysis is a newly developed mathematical framework that enables this integration and allows for analysis of multimodal data [15, 24, 25]. A multilayer network is a ‘network of networks’, comprised of multiple interconnected layers, each characterizing a different aspect of the same system. Figure 1 illustrates the concept of multilayer networks using the analogy of a commuter network. Although the framework of multilayer networks is relatively new in the field of neuroscience, promising results have already been reported. Multilayer analysis of dMRI and fMRI networks of healthy participants confirmed the synergistic nature of the structure and function of the brain network [26]. Further relevance of multilayer analysis has been shown in clinical studies: multilayer connectivity differences were reported between patients with schizophrenia and healthy controls, and these differences were related to symptom severity [27]. Moreover, a study in schizophrenia and another study in Alzheimer’s disease suggested that multilayer centrality could outperform unilayer measures to distinguish cases from healthy controls [28, 29]. Additionally, an MEG study used nodal centrality metrics to identify brain regions that were vulnerable in patients with Alzheimer’s disease compared to healthy controls, and found that such regions could only be detected using a multilayer approach. Even more relevant to our work, this vulnerability of central regions in the multilayer network was related to cognitive dysfunction [30]. Multilayer network analysis can thus contribute to a better understanding of the relation between the FPN and EF.

**Figure 1.**
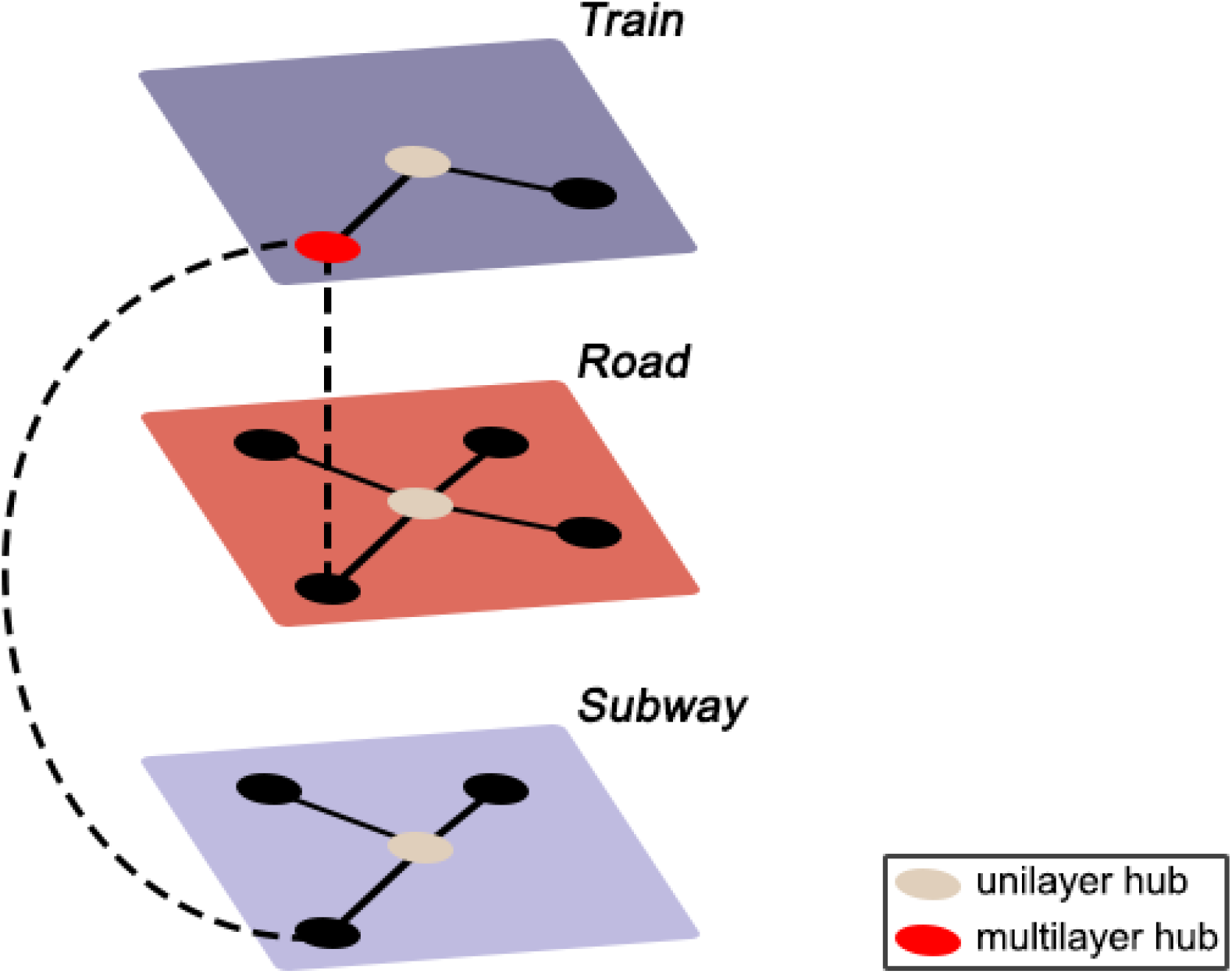
Example of a multilayer transport network, encoding information about train, subway, and road connectivity, and the interlayer links between them. Suppose there is a sudden increase in commuters in the subway network in the absence of any delayed or cancelled subway cars or suspended subway stations. Considering the unilayer subway network in isolation, the observed spike in commuters would seem inexplicable, as the properties of the subway network (i.e. the links and nodes) are unaltered. However, observing the entire transport system might reveal severe delays in the train network, forcing people who usually commute by train to now use the subway, thus leading to an increase in commuters in the subway network. Likewise, consider the red node in the train network. From a unilayer perspective, this is a peripheral station that is of little importance to the transport system. However, the multilayer perspective reveals this to be the only location where all three modes of transport connect, and the seemingly peripheral station thus plays a significant integrative role in the transportation network – a property that would have remained unnoticed without incorporating all the layers of the system.

Here, we used multimodal data to assess the association between FPN centrality and EF in healthy participants and explored the potential added value of a multilayer framework over a unilayer framework. We hypothesized 1) a positive association between FPN centrality of both the uni- and multilayer networks and EF, and 2) multilayer centrality superseding its unilayer equivalents in explaining individual differences in EF.

## 2. Methods

### 2.1. Participants

This study was preregistered in the Netherlands Trial Register under trial ID NL7301. Thirty-nine (39) healthy participants were prospectively recruited for this specific study through an online platform, Hersenonderzoek.nl (www.hersenonderzoek.nl), where volunteers can register for participation in neuroscience studies. Participants were selected based on the following inclusion criteria: (1) age between 20 and 70 years old; (2) native Dutch speaker; (3) able to provide written informed consent. The following exclusion criteria were used: (1) history of neurological or psychiatric disease; (2) current and regular use of centrally acting drugs; (3) presence of contraindications for MRI or MEG. Participants were asked not to ingest any caffeine or alcohol on the testing days. Approval was obtained from the VU University Medical Center Medical Ethical Committee, and all subjects provided written informed consent prior to participation.

### 2.2. Neuropsychological evaluation

Participants underwent an extensive customized neuropsychological test battery, consisting of the Dutch version of Rey’s Auditory Verbal Learning Test [31], the Concept Shifting Test (CST; [32]), the Memory Comparison Test (MCT), the Stroop Color-Word Test (SCWT; [33]), the Location Learning Test (LLT; [34]), the Categorical Word Fluency Test [35], and the Letter-Digit Modalities Test (LDMT; [36]). We used (subscores on) three of these tests to assess EF. The first test we used was the CST, where the participant was shown 16 small circles, grouped in a large circle, containing either digits (CST part A), letters (CST part B), or both digits and letters (CST part C). These circles needed to be crossed out in ascending order in part A, in alphabetical order in part B, and in alternating order (digit-letter) in part C. The participant was asked to perform the test as quickly as possible without making mistakes. Additionally, to correct for motor speed, a null-condition with empty circles (CST zero) was carried out thrice. The second test we used was the SCWT, where the participant was asked to read four different cards. On the first card, names of colors – red, green, yellow, and blue – were printed in black ink. On the second card, rectangles were printed in these same colors. On the third card, the names of the colors were printed in an inconsistent color ink; e.g. the word ‘red’ was printed in yellow ink, and the participant was asked to read the color of the ink and ignore the word. The fourth card was identical to the third card, but several words were circled. For these circled words, the participant was asked to read the word itself instead of the color of the ink. The third test we used was the Word Fluency Test, where the participant was asked to name as many words in the category ‘animals’ as possible within 60 seconds.

Using validated norms of the CST [32], SCWT [37], and Word Fluency Test [37], raw scores were adjusted for sex, age, and education (classified according to the Dutch Verhage system [38] which ranges from level 1 [less than six years of primary education] to level 7 [university degree]) and transformed into z-scores. EF was defined as the average of z-scores for Word Fluency, Stroop-interference (time to complete card 3 corrected for the time to complete card 2), and CST-shift (time to complete card C minus the average time to complete cards A and B, adjusted for time to complete CST zero) [32].

### 2.3. Magnetic resonance imaging

MRI data were obtained using a 3T MRI system (Philips Ingenia CX) with a 32-channel receive-only head coil at the Spinoza Centre for Neuroimaging in Amsterdam, The Netherlands. A high-resolution 3D T1-weighted image was collected with a magnetization-prepared rapid acquisition with gradient echo (MPRAGE; TR = 8.1ms, TE = 3.7ms, flip angle = 8°, voxel dimensions = 1 mm3 isotropic). This anatomical scan was registered to MNI space through linear registration with nearest-neighbor interpolation, and was used for coregistration and normalization of all other modalities (dMRI, fMRI, and MEG) to the same space.

#### 2.3.1. Diffusion MRI

Diffusion MRI was collected with diffusion weightings of b = 1000 and 2000 s/mm^2^ applied in 29 and 59 directions, respectively, along with 9 non-diffusion weighted (b = 0 s/mm^2^) volumes using a multiband sequence (MultiBand SENSE factor = 2, TR = 4.7 s, TE = 95 ms, flip angle = 90°, voxel dimensions = 2 mm^3^ isotropic, no interslice gap). In addition, two scans with opposite phase encoding directions were collected for blip-up blip-down distortion correction using FSL topup [39]. Structural connectomes were constructed by performing probabilistic Anatomically-Constrained Tractography (ACT) [40] in MRtrix3 [41]. A tissue response function was estimated from the pre-processed and bias field corrected dMRI data using the multi-shell multi-tissue five-tissue-type algorithm (msmt_5tt). Subsequently, the Fiber Orientation Distribution (FOD) for each voxel was determined by performing Multi-Shell Multi-Tissue Constrained Spherical Deconvolution (MSMT-CSD) [42]. ACT was performed by randomly seeding 100 million fibers within the white matter to construct a tractogram, and Spherical-deconvolution Informed Filtering of Tractograms (SIFT, SIFT2 method in MRtrix3) [43] was then performed to improve the accuracy of the reconstructed streamlines and reduce false positives. For every participant, their respective 3D T1-weighted image was used to parcellate the brain into 210 cortical Brainnetome atlas (BNA) [44] regions. We then used this parcellation to convert the tractogram to a structural network, where weighted edges represented the sum of all streamlines leading to and from all voxels within two brain regions.

#### 2.3.2. Resting-state functional MRI

Resting-state fMRI was collected using a multiband sequence (MultiBand SENSE factor = 2, TR = 1.52 s, TE = 30 ms, flip angle = 70°, voxel size = 2.5 × 2.5 × 2.75 mm^3^, interslice gap = 0.25 mm, 310 volumes, 12-min acquisition). Participants were instructed to remain awake with their eyes open. Pre-processing was done using FSL 5 (FMRIB 2012, Oxford, United Kingdom, http://www.fmrib.ox.ac.uk/fsl) and included brain extraction, removal of the first four volumes, motion correction by regressing out six motion parameters, and spatial smoothing at 5 mm full-width-half-maximum (FWHM). An independent component analysis was performed for Automatic Removal of Motion Artefacts (ICA-AROMA) [45], followed by regressing out white matter and cerebrospinal fluid signals and high-pass filtering (100 s cutoff). Mean absolute motion did not exceed 0.6 mm for any participant; the median was 0.27 mm (0.08-0.59 mm). The rsfMRI data were registered to native 3D T1 space using boundary-based registration. The BNA atlas was then reverse-registered to each participant’s functional data using nearest-neighbor interpolation. For every participant, a mask containing only grey matter voxels with reliable rsfMRI signal was constructed by combining a grey matter mask and an rsfMRI mask, excluding all voxels with a signal intensity in the lowest quartile of the robust range (see [46] for more details). Time-series were extracted from all atlas regions by averaging time-series across all voxels within each region. Thirteen regions with signal loss (i.e. regions with zeros in the functional connectivity matrices) due to magnetic field inhomogeneities in these echo-planar imaging (EPI) sequences were removed from further analyses across all participants and modalities. Thus, 197 atlas regions remained for all further analyses. Finally, for every participant, Pearson correlation coefficients between all pairs of time-series were calculated to obtain a functional connectivity matrix. Correlation coefficients were absolutized, as most network metrics do not take into account negative values, but inverse correlations may carry relevant information [47, 48].

### 2.4. Magnetoencephalography

MEG data were recorded in a magnetically shielded room (Vacuumschmelze GmbH, Hanau, Germany) using a 306-channel (102 magnetometers and 204 gradiometers) whole-head MEG system (Elekta Neuromag Oy, Helsinki, Finland) with a sampling frequency of 1250 Hz during a no-task, eyes-closed condition for five minutes, an eyes-open condition for two minutes, and a final eyes-closed condition for another five minutes, with the participant in supine position. Here, we used only the first eyes-closed recording for all further analyses. An anti-aliasing filter of 410 Hz and a high-pass filter of 0.1 Hz were applied online. The cross-validation Signal Space Separation (xSSS) [49] was applied to aid visual inspection of the data. We removed channels containing no signal or noisy signal, with a maximum of 12 channels removed per participant. Further noise removal was performed offline using the temporal extension of Signal Space Separation (tSSS) [50] in MaxFilter (version 2.2.15). The head position relative to the MEG sensors was recorded continuously using the signals from five head-localization coils. Coil positions and the scalp outline were digitized using a 3D digitizer (Fastrak, Polhemus, Colchester, VT, USA). A surface-matching procedure was used to achieve co-registration of the participant’s digitized scalp surface and their anatomical MRI, with an estimated resulting accuracy of 4 mm [51]. A single best-fitting sphere was fitted to the outline of the scalp as obtained from the co-registered MRI, which was used as a volume conductor model for the beamformer approach described below. The co-registered MRI was spatially normalized to a template MRI, and the voxels in the normalized co-registered MRI were again labeled according to the same atlas. We then used a scalar beamforming approach [52] to reconstruct the source of neurophysiological activity from the sensor signal. The beamformer weights were based on the lead fields, the broadband (0.5-48 Hz) data covariance, and noise covariance. The data covariance was based on, on average, 298 s of data (range 293-314 s). A unity matrix was used noise covariance. Broadband data were then projected through the normalized beamformer weights to obtain time-series for each atlas region. Out of all the voxels that constitute an atlas region, the centroid [53] was selected to reconstruct localized MEG activity, resulting in time-series for each of the 197 included cortical regions. For all participants, we included the first 88 epochs of 4096 samples (3.28s) of the obtained time-series (total length 4 minutes and ~48 seconds). Fast Fourier transforms were applied to filter the time-series into six frequency bands: delta (0.5-4 Hz), theta (4-8 Hz), lower alpha (8-10 Hz), upper alpha (10-13 Hz), beta (13-30 Hz), and gamma (30-48 Hz). We then computed the phase lag index (PLI) [54] between the frequency-filtered time-series of all pairs of regions using custom-made scripts in MATLAB (R2018b, Mathworks, Natick, MA, USA) to obtain weighted functional connectivity matrices.

### 2.5. Unilayer network construction and analysis

First, we constructed minimum spanning trees (MST) for the six frequency-band specific MEG networks by applying Kruskal’s algorithm [55] to the functional connectivity matrices. The MST is a binarized sub-graph of the original graph that connects all the nodes in the network without forming loops. This represents the backbone of the network [56, 57] and, importantly, is not hindered by common methodological issues such as effects of connection strength or link density on the estimated topological characteristics of networks [57]. Edge weights were defined as the inverted PLI values (1/PLI) when constructing the *minimum* spanning tree, since we were interested in the strongest connections [58].

We then calculated nodal eigenvector centrality (EC) individually for each of the six MEG MSTs, and for the fully connected weighted dMRI and rsfMRI connectivity matrices, using the brain connectivity toolbox (https://sites.google.com/site/bctnet/) in MATLAB. EC is a measure of nodal centrality that assumes that a node is more influential if it is connected to nodes that are highly central themselves, and thus considers both the connections of a node itself as well as the connections of its neighbors. This makes it an interesting measure of centrality that takes the entire network into account, and it has been shown to be highly relevant for cognition in studies using dMRI [59], rsfMRI [46], and MEG [60]. For a more detailed explanation of the EC and its mathematical definition, see [61].

Finally, we extracted and subsequently averaged the ECs of all nodes belonging to the FPN to obtain one value per unilayer network per participant (for a total of eight values per participant). Regions belonging to the FPN were defined based on an earlier categorization [62] of the regions of the BNA according to the classical seven-network parcellation by Yeo and colleagues [63].

### 2.6. Multilayer network construction and analysis

A multiplex network is a multilayer network used to describe different interactions between the same set of nodes [64]. In this context, each layer is characterized by a different modality of interaction. Therefore, this mathematical framework is useful to encode information from brain networks created using different edge weights or imaging modalities as long as all layers are built using the same atlas. In such a multiplex network, links between different layers, also known as interlayer links, exclusively connect the same node or brain region across layers.

There is, as of yet, no established method for determining biologically meaningful weighted interlayer links between different modalities. Additionally, network metrics can potentially be biased by differences in link density and average connectivity across layers and between participants [23]. Here, we therefore decided to construct binary multiplex networks. Consequently, in addition to the MEG MSTs described in section 2.5, we used Kruskal’s algorithm to construct MSTs for the dMRI and rsfMRI data. We then integrated these eight MSTs to obtain an interconnected multiplex network for every participant. Each participant’s multiplex thus consisted of L = 8 layers (one for dMRI, one for rsfMRI, and one for each of the six MEG frequency bands), with each layer containing the same set of N = 197 nodes (atlas regions), and each spanning tree and thus layer having M = N – 1 = 196 intralayer links. The weights of the interlayer connections were set to 1, identical to the intralayer connections. The resulting multilayer network was represented as an LxN by LxN supra-adjacency matrix (see Figure 2) with diagonal blocks encoding intralayer connectivity for each modality and off-diagonal blocks encoding interlayer connectivity. Supra-adjacency matrices were then exported to Python (version 3.6, Python Software Foundation, available at http://www.python.org), and multilayer nodal EC (see [64] for a mathematical definition) was computed using custom-made scripts that integrate the Python libraries multiNetX [65] and NetworkX (version 2.3) [66] that can be found on GitHub (https://github.com/nkoub/multinetx and https://github.com/networkx, respectively). Note that EC was first computed for each node in each layer separately and subsequently aggregated across layers to obtain one value per node, as described earlier [67]. We then again extracted and averaged ECs of the FPN nodes, yielding one value for multilayer EC per participant. A schematic overview of the methods can be found in Figure 2; Figure 3 shows an example multiplex network as constructed using these methods. All of the custom-made scripts, as well as the data that we used in this study, can be found on our lab’s GitHub page (https://github.com/multinetlab-amsterdam/projects/tree/master/mumo).

**Figure 2.**
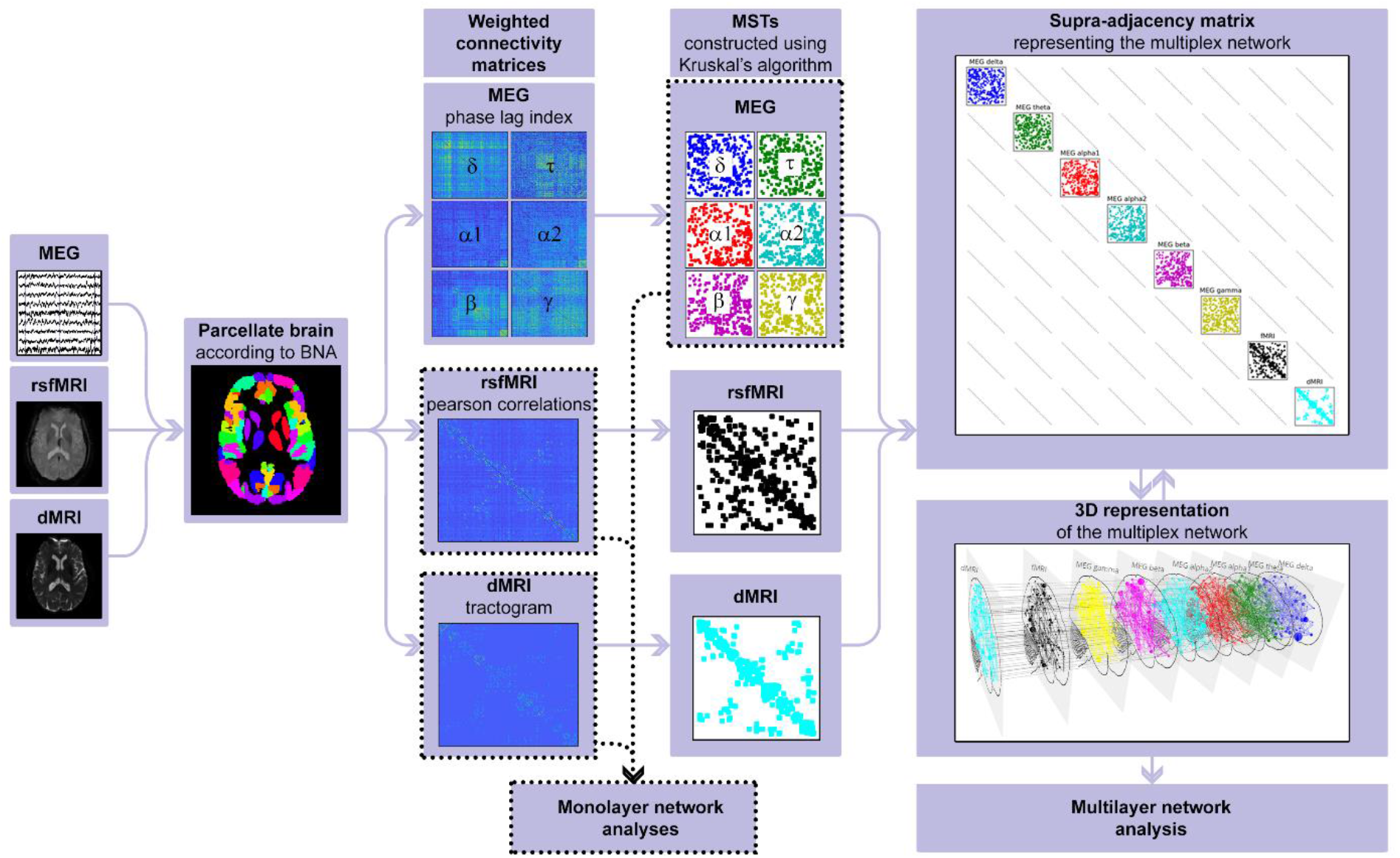
Schematic overview of the analysis pipeline. For every participant, raw imaging data obtained from diffusion MRI, resting-state functional MRI, and magnetoencephalography was pre-processed; the brain was parcellated according to the brainnetome atlas; connectivity was calculated to construct weighted connectivity matrices; minimum spanning trees of the weighted matrices were constructed using Kruskal’s algorithm; and finally a supra-adjacency matrix representing a multilayer network was constructed. Note that unilayer network measures were computed on the minimum spanning trees of the magnetoencephalography frequency bands, but weighted data was used for diffusion MRI and resting-state functional MRI. MEG = magnetoencephalography. rsfMRI = resting-state functional MRI. dMRI = diffusion MRI. BNA = brainnetome. MST = minimum spanning tree.

**Figure 3.**
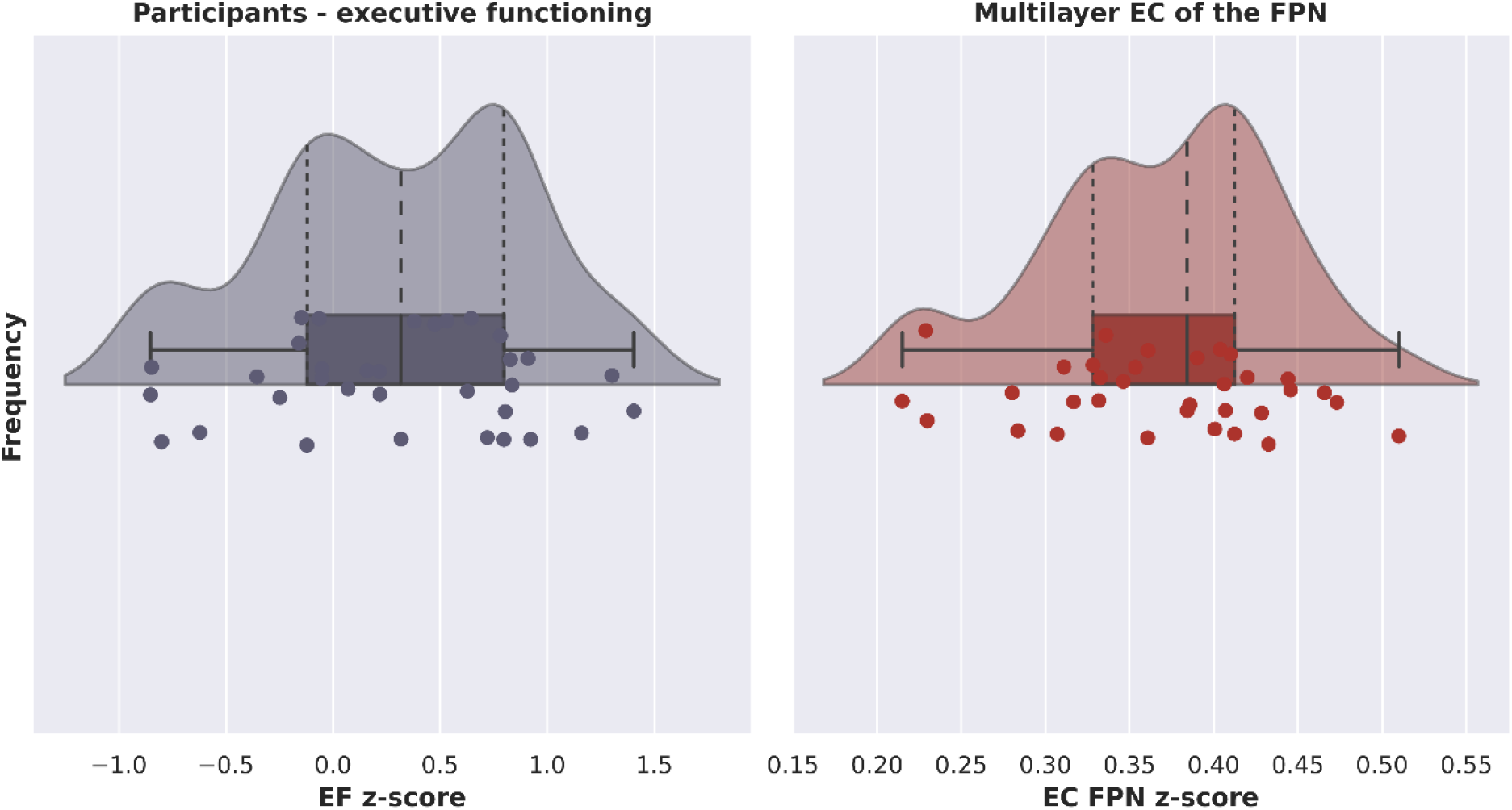
Raincloud plots showing probability density, summary statistics, and individual datapoints of the z-score per participant of executive functioning (left) and multilayer eigenvector centrality of the frontoparietal network (right). EF = executive functioning. EC = eigenvector centrality. FPN = frontoparietal network.

### 2.7. Statistical analyses

To assess the relation between uni- or multilayer EC of the FPN and age, sex and education-corrected EF scores, we performed a multiple regression analysis in SPSS (version 26, IBM Corp., Armonk, NY, USA). With EF as the dependent variable, average EC values of the FPN of each of the eight unilayer networks described in section 2.5 were added in a first block using a backward stepwise procedure (F probability for removal 0.10), and the average EC of the FPN of the multilayer network was entered in a second block. To assess whether these data met the assumption of collinearity, we computed bivariate correlations between all average unilayer EC values of the FPN, and additionally ran collinearity diagnostics.

## 3. Results

### 3.1. Participant characteristics

Of the 39 included participants, two participants dropped out before completion of the study, two were excluded during the study because of contra-indications for MRI, and another two were excluded after visual inspection of their MRI data revealed artifacts. This resulted in a total of 33 included participants with complete structural MRI, dMRI, rsfMRI, MEG, and neuropsychological data that were used in the analyses. Of these participants, 18 were female and 15 were male. They were well spread out in terms of age, ranging between 22 and 70 years old, with a mean age of 46 ± 17 years. Participants were mainly higher-educated.

### 3.2. Network correlates of executive functioning

Figure 4 shows a raincloud plot with the distribution of EF z-scores for all participants. Importantly, as indicated by the wide range of normed z-scores, our sample was diverse in terms of EF performance. There was no evidence of multicollinearity between network variables: absolute correlation coefficients between the unilayer network eigenvector centralities did not exceed 0.7, and tolerance values were all greater than 0.1. Figure 5 shows, for one participant, exemplar values of EC for the multilayer network, as well as all the unilayer networks and the mean of the unilayers.

**Figure 4.**
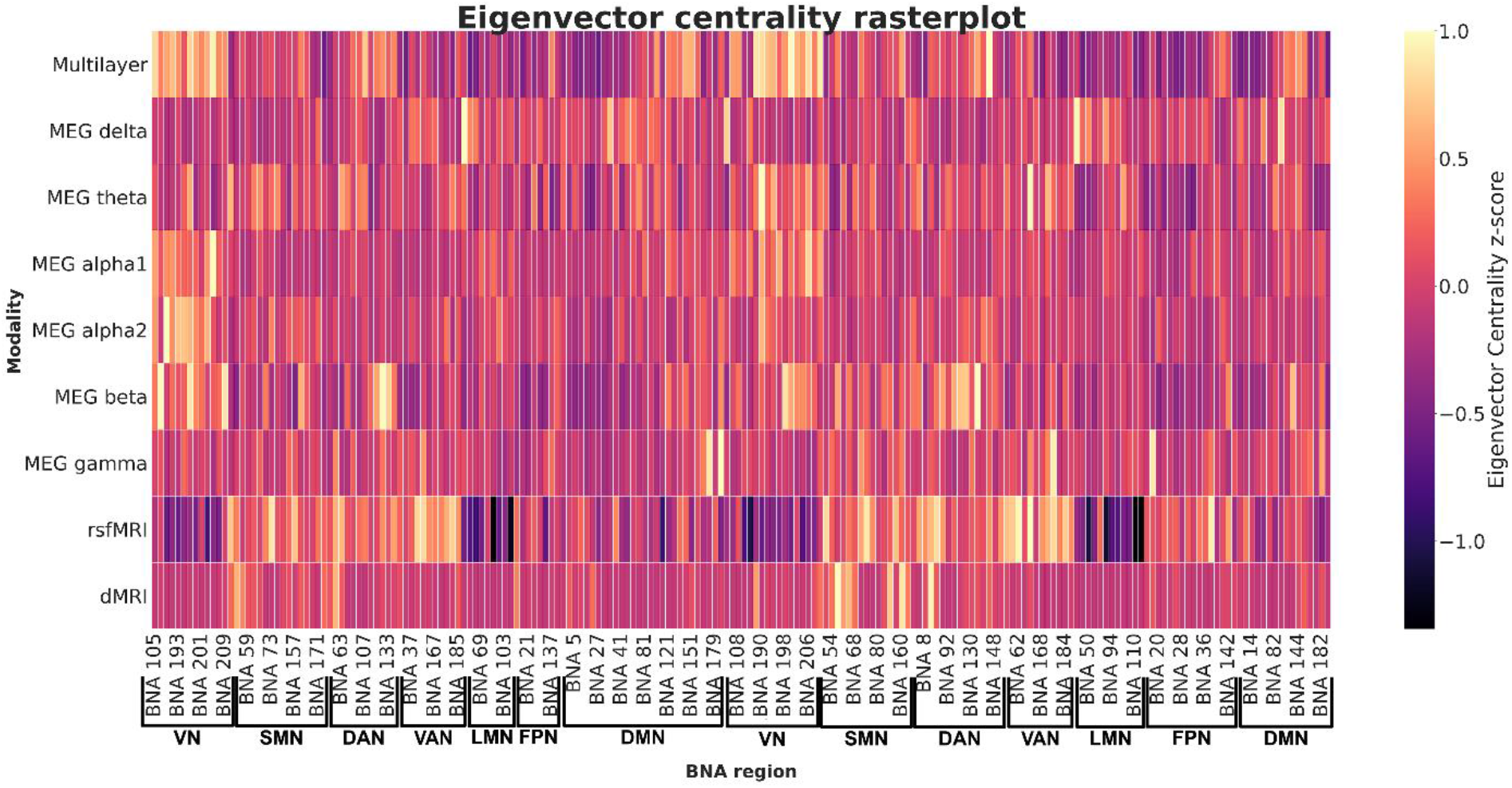
Rasterplot showing multilayer eigenvector centrality in the eight unilayer networks and the multilayer network, ordered by subnetwork (all left-hemisphere regions followed by all right-hemisphere regions). Yellow indicates regions with high EC. This shows the differences in ‘centrality profiles’ across modalities. BNA region numbers refer to the labels as given in Supplementary Table 1. MEG = magnetoencephalography. rsfMRI = resting-state functional MRI. dMRI = diffusion MRI. BNA = brainnetome atlas. VN = visual network. SMN = somatomotor network. DAN = dorsal attention network. VAN = ventral attention network. LMN = limbic network. FPN = frontoparietal network. DMN = default mode network.

**Figure 5.**
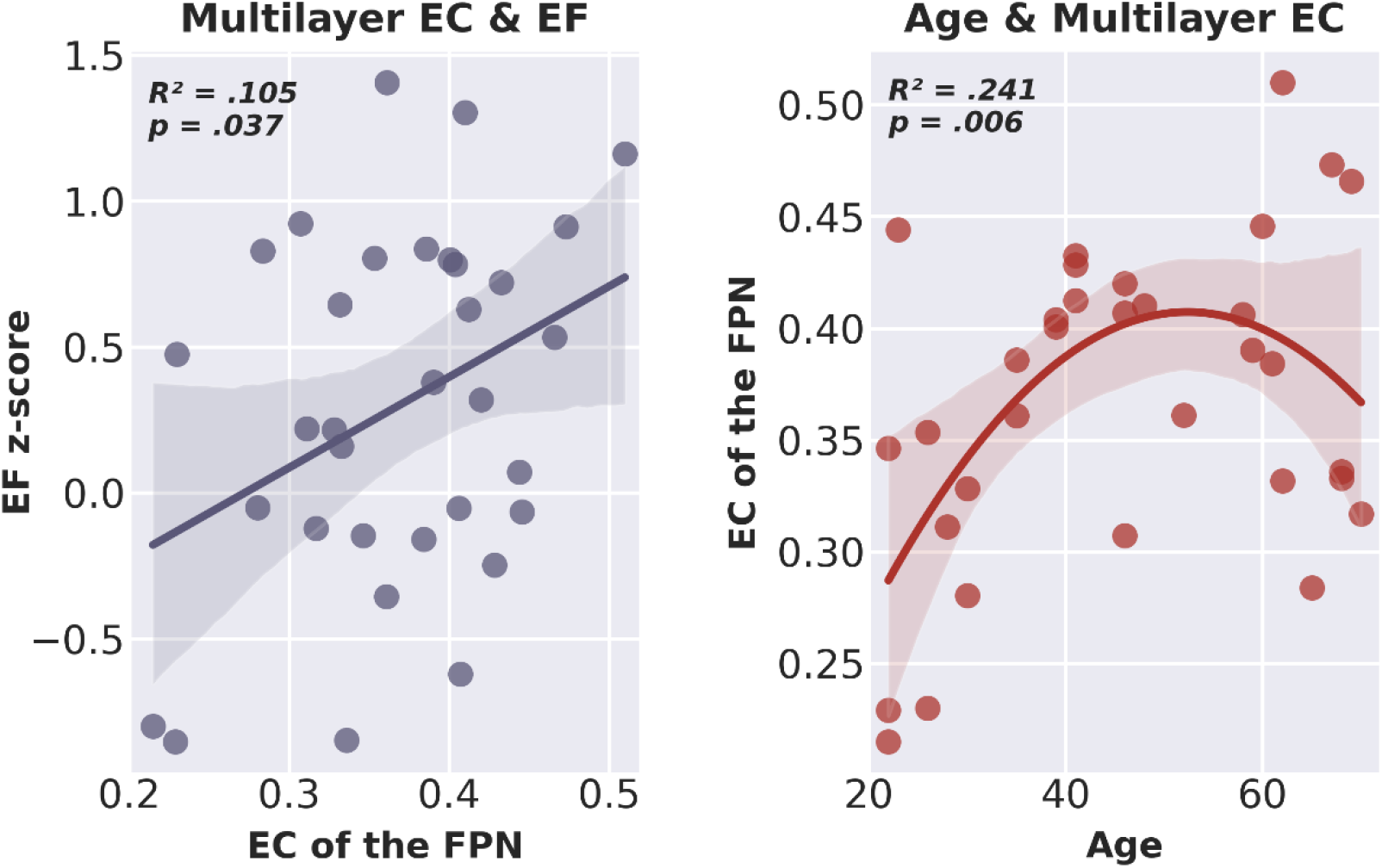
Scatterplot including line of best fit of multilayer eigenvector centrality of the frontoparietal network and executive functioning (left) and age and multilayer eigenvector centrality of the frontoparietal network (right). EC = eigenvector centrality. EF = executive functioning. FPN = frontoparietal network.

Testing our hypotheses, none of the unilayer network eigenvector centralities survived the backwards stepwise selection, see Table 1 for the coefficients of the included and excluded variables. The final regression model, containing only multilayer EC of the FPN as a predictor of EF, was statistically significant (*R*^2^ =.133, adjusted *R*^2^ = .105, *F*[1, 31] = 4.753, *p* = .037). There was no significant increase in *R*^2^ from the second-to-last model, containing two predictors (unilayer EC in the lower alpha band and multilayer EC), to this final significant model. These results suggest that only EC of the FPN of the multilayer network was a significant predictor of EF, and that a higher multilayer EC of the FPN was related to better EF (see Figure 6).

**Table 1.** Standardized Beta coefficients p-values of included and excluded variables of the regression models. Top: multilayer eigenvector centrality of the frontoparietal network and executive functioning. Bottom: age and multilayer eigenvector centrality of the frontoparietal network. EC = eigenvector centrality. FPN = frontoparietal network. EF = executive functioning. MEG = magnetoencephalography. dMRI = diffusion MRI. rsfMRI = resting-state functional MRI. * indicates significance at the p<0.05 level.

| **Multilayer EC of the FPN & EF** |  |  |
| --- | --- | --- |
|  | **$\beta$** | ***P*** |
| **Final model ($R^2_{adj} = .105$)** |  |  |
| Multilayer EC | .365 | .037* |
| **Excluded variables** |  |  |
| EC MEG delta | .047 | .798 |
| EC MEG upper alpha | .056 | .752 |
| EC dMRI | .074 | .669 |
| EC MEG theta | -.065 | .716 |
| EC rsfMRI | .097 | .587 |
| EC MEG beta | .144 | .407 |
| EC MEG gamma | -.204 | .234 |
| EC MEG lower alpha | -.238 | .163 |
| **Age & multilayer EC of the FPN** |  |  |
|  | **$\beta$** | ***P*** |
| **Final model ($R^2_{adj} = .241$)** |  |  |
| Age | .014 | .010* |
| Age squared | .00013 | .021* |

Repeating these analyses using a forward stepwise procedure (F probability for entry 0.05) resulted in an identical final model, containing only multilayer EC as a predictor of EF and was significant (*R*^2^ = .133, adjusted *R*^2^ =.105, *F*[1, 31] = 4.753, *p* = .037).

### 3.3. Multilayer network correlates of age

To further validate the relevance of multilayer FPN centrality, we tested its relationship with age. Unilayer network studies have revealed that the brain network tends to become more efficiently integrated in early life [68], after which its development plateaus during middle age [69] and subsequently regresses to a less integrative topology with older age [70]. This relation is also reflected in changes in for example whole brain- [71] and white matter volume [72, 73] across the lifespan, and the effects of age on the development of neurological disease, e.g. in multiple sclerosis [74] or Alzheimer’s disease [75], is well-established. We therefore hypothesized an inverted-U relation between age and multilayer EC of the FPN. We employed a hierarchical multiple regression model with multilayer centrality as the dependent variable. Age was entered in a first block, and the square of age was added to the model in a second block. We used an alpha level of .05 for all statistical tests. Both regression models were checked for normality of residuals using a Q-Q plot.

See figure 4 for a raincloud plot of multilayer centrality, showing the distribution of multilayer network EC of the FPN for all participants. The final model with both age and age squared indicated a statistically significant quadratic relation between age and multilayer EC of the FPN (*R*^2^ = .289, adjusted *R*^2^ = .241, *F*[2, 30] = 6.082, *p* = .006). The square of age added significantly to the model, leading to an increase in *R*^2^ of .140 (F[1, 30] = 5.915, *p* = .021), suggesting indeed that the quadratic model more accurately explained age variations than the simple linear model. The coefficients of the included variables are reported in Table 1; Figure 6 shows the relation between age and multilayer EC of the FPN.

## 4. Discussion

We studied how multilayer centrality of the FPN was related to individual differences in EF, and whether this provided additional information to modality- and frequency-specific unilayer FPN centrality. We found that higher multilayer FPN centrality related to better EF, whereas FPN centrality of unilayer networks did not significantly explain differences in EF between healthy adults. Finally, *post hoc* analyses established an inverted-U relationship between age and multilayer centrality of the FPN.

Firstly, at least for the multilayer network, these results are in line with other studies displaying the importance of FPN network centrality for EF. The relation between FPN centrality and, by extension, network integration and cognition has been well-established in unilayer networks, using different neuroimaging and neurophysiological modalities. Increased integration of the FPN within the entire brain network specifically has been related to better EF in studies utilizing dMRI [11], rsfMRI [12, 13], and MEG [14]. While network segregation is thought to enable fast processing of lower-order information (e.g. analysis of visual inputs) [76], highly central nodes like those within the FPN facilitate global communication between these segregated communities, presumably enabling higher-order cognitive processes and specifically EF (see e.g. [9, 77]).

Moreover, our results demonstrate the relevance of multimodal network analysis through a multilayer network approach in explaining cognitive variance. While FPN centrality of the unimodal networks did not relate significantly to EF, higher FPN centrality of the multilayer networks was indeed associated with better EF. Visual exploration of our data confirmed that the level of integration per node depends on the modality on which the network is based, and that this is again different for the multilayer network. Central nodes (i.e. nodes with high EC) in the multilayer network are thus not the same as central nodes in the monolayer networks (see Figure 5). Other multilayer studies have similarly reported that the precise node that can be considered most central in an unilayer network, may not serve as the most central node in a multilayer network and vice versa [29, 30]. We build upon these studies by demonstrating that multimodal information captures variance in EF that networks obtained from a single modality do not.

Finally, the quadratic relation between age and multilayer centrality possibly reflects the rise and decline of brain network efficiency across the life span, and is in line with findings from studies reporting on unimodal data. Unilayer brain networks have been shown to become more segregated or modular during development [78], and connectivity of highly central regions has been reported to increase from childhood to adulthood [68], suggesting that the brain network becomes increasingly efficient with maturation. However, after a certain age, modularity of the network seems to decrease [70], indicating a degradation of the efficiency of the brain network. Moreover, a similar plateauing of brain network efficiency around middle age has been reported in the organization of the ‘rich club’ [69]. We have shown here that this characteristic of the brain network is maintained in a multilayer network, showing that FPN centrality of the multilayer network is an age-relevant metric. Note that we corrected EF scores for age, such that the association we find between EF and multilayer centrality cannot be ascribed to age effects alone. However, larger datasets are needed to disentangle the exact relationship between age, network centrality, and EF.

The biological interpretation of the multilayer network used in this work deserves further consideration. Importantly, the spatial definition of nodes is identical across layers: the nodes in each modality were defined based on the same brain regions. The use of the brainnetome atlas, which is based on both structural- and functional connectivity pattern similarity within and across brain regions, further supports the assumption that these nodes can indeed be seen as canonical units across layers. We then used interlayer links between the same brain regions (nodes) across layers to integrate different modalities. The biological assumption here is that structure and function conflate maximally within the same brain region. There is ample evidence that this assumption holds across macroscopic modalities when correlating, for instance, structural and rsfMRI connectivity patterns across the whole brain [20, 22, 79]. The spatial variation that exists in nodal correlations between structural and functional connectivity [80], however, may indicate that although this connectivity is highest within the same region instead of between regions, the linkage between layers varies per region. Such variations were not taken into account in the current work, where we used MSTs of the individual layers for the construction of multiplex networks and set the weights of all interlayer connections in the multilayer networks to one. Future studies may therefore incorporate weighted interlayer links to represent the spatial variation in within-region correlations across modalities. Another potential shortcoming of the binarization of link weights is that it eliminates layer dominance [81]: some layers may have a stronger influence on multilayer network characteristics than others, but when all layers carry the same importance, this information is lost. However, just as some layers may drive the properties of the multilayer more strongly than other layers, other layers may play a negligible role. This raises the question whether all possible layers should be included, or whether an a priori selection should be made – and if so, how this selection should be made. A first foray into this issue was made in a dataset of social contacts [82], but the layer selection problem in multimodal brain networks warrants further exploration. Furthermore, although interlayer connectivity may be maximal within brain regions, there is potential connectivity between different regions across different modalities-- i.e., cross-talk between node A in modality × and node B in modality Y may be relevant to overall functioning of the network. A general multilayer network formulation allows interlayer links between all nodes in all layers (see e.g. [64]). However, multimodal datasets present considerable challenges when constructing a full multilayer network. Chief among them is determining biologically meaningful interlayer links between different modalities at the individual participant level. Additionally, a recent modelling study revealed that interlayer connectivity is driven mostly by one-to-one (i.e. multiplex) connections [83], and as evidenced by previous empirical studies [29, 30], a multiplex approach is therefore a logical and intuitive first step for analyzing multidimensional data. Lastly, we defined the FPN based on a predefined classification. As such, the FPN was comprised of the same regions across the different unilayers as well as in the multilayer network. However, subnetworks like the FPN and the hubs within them have been found to vary depending on the modality used [22, 84], and have also been shown to be different in a multiplex compared to a unilayer network [29]. Additionally, there is a large individual variability in the functional topography of the FPN [85]. A more data-driven approach to the formulation of the FPN may therefore further increase the explanatory power of the multilayer approach.

Some additional limitations need to be taken into consideration. It is particularly important to take into account the relatively small sample size of the present study: the absence of any unilayer effects could be due to a lack of statistical power, rather than the true absence of any correlations between unilayer network metrics and cognition. Potentially related to this limited sample size, the model containing only the significant multilayer predictor was not significantly better than the model containing a nonsignificant unilayer plus the multilayer predictor in terms of its explanatory value. Despite the significant association between multilayer FPN centrality and EF, we therefore cannot conclude with certainty that multilayer FPN centrality is more valuable than unilayer FPN centrality in explaining EF based on this study. Also, for the unilayer analyses, we computed network metrics based on the MSTs of the MEG networks but utilized the fully connected weighted networks of the dMRI and rsfMRI data. We made this choice to conform with previous modality-specific literature (e.g. [58, 86, 87]).

## 5. Conclusions

Integration of multimodal brain networks through a multilayer framework relates to EF in healthy adults, and corroborates known brain associations with aging. These findings underline the relevance of a multimodal view on integration of the brain network as a correlate of EF. Furthermore, the multilayer approach may be of particular interest in populations where network alterations differ across modalities. For instance, early neurodegeneration and structural network deterioration of particularly the most central regions in the brain are initially associated with increases in functional communication, after which the functional brain networks seems to collapse [88, 89]. Multilayer network analysis may advance our understanding of the interplay between structural and different functional network aspects in such clinical populations.

## Acknowledgements

The authors would like to thank the Amsterdam Neuroscience research institute for supporting this study. We would also like to thank Hersenonderzoek.nl, a Dutch online registry that facilitates participant recruitment for neuroscience studies (www.hersenonderzoek.nl). Hersenonderzoek.nl is funded by ZonMw-Memorabel (project no 73305095003), a project in the context of the Dutch Deltaplan Dementie, Gieskes-Strijbis Foundation, the Alzheimer’s Society in the Netherlands and Brain Foundation Netherlands. We thank the laboratory technicians of the Amsterdam UMC, Department of Clinical Neurophysiology and MEG Center, as well as the scan assistants of the Spinoza Centre for Neuroimaging, for the data acquisition. Finally, we thank all participants for their participation.

