## Supplemental Table 1 for "Multimodal multilayer network centrality relates to executive functioning"

Supplementary Table 1

*Brainnetome atlas regions including anatomical label and subnetwork allocation. The following regions were excluded from analyses and are crossed out: 47, 48, 69, 70, 94, 101, 111, 113, 115-119. VAN = ventral attention network. DMN = default mode network. DAN = dorsal attention network. SMN = somatomotor network. FPN = frontoparietal network. LMN = LMN network. VN = VN network.*

| **BNA region** | **BNA label** | **Anatomical description** | **Subnetwork** |
| --- | --- | --- | --- |
| 1 | Left_A8m | medial area 8 | VAN |
| 2 | Right_A8m | medial area 8 | VAN |
| 3 | Left_A8dl | dorsolateral area 8 | DMN |
| 4 | Right_A8dl | dorsolateral area 8 | DMN |
| 5 | Left_A9l | lateral area 9 | DMN |
| 6 | Right_A9l | lateral area 9 | DMN |
| 7 | Left_A6dl | dorsolateral area 6 | DAN |
| 8 | Right_A6dl | dorsolateral area 6 | DAN |
| 9 | Left_A6m | medial area 6 | SMN |
| 10 | Right_A6m | medial area 6 | SMN |
| 11 | Left_A9m | medial area 9 | DMN |
| 12 | Right_A9m | medial area 9 | DMN |
| 13 | Left_A10m | medial area 10 | DMN |
| 14 | Right_A10m | medial area 10 | DMN |
| 15 | Left_A9/46d | dorsal area 9/46 | VAN |
| 16 | Right_A9/46d | dorsal area 9/46 | FPN |
| 17 | Left_IFJ | inferior frontal junction | FPN |
| 18 | Right_IFJ | inferior frontal junction | FPN |
| 19 | Left_A46 | area 46 | FPN |
| 20 | Right_A46 | area 46 | FPN |
| 21 | Left_A9/46v | ventral area 9/46 | FPN |
| 22 | Right_A9/46v | ventral area 9/46 | FPN |
| 23 | Left_A8vl | ventrolateral area 8 | DMN |
| 24 | Right_A8vl | ventrolateral area 8 | FPN |
| 25 | Left_A6vl | ventrolateral area 6 | DAN |
| 26 | Right_A6vl | ventrolateral area 6 | FPN |
| 27 | Left_A10l | lateral area10 | DMN |
| 28 | Right_A10l | lateral area10 | FPN |
| 29 | Left_A44d | dorsal area 44 | FPN |
| 30 | Right_A44d | dorsal area 44 | FPN |
| 31 | Left_IFS | inferior frontal sulcus | FPN |
| 32 | Right_IFS | inferior frontal sulcus | FPN |
| 33 | Left_A45c | caudal area 45 | DMN |
| 34 | Right_A45c | caudal area 45 | FPN |
| 35 | Left_A45r | rostral area 45 | DMN |
| 36 | Right_A45r | rostral area 45 | FPN |
| 37 | Left_A44op | opercular area 44 | VAN |
| 38 | Right_A44op | opercular area 44 | VAN |
| 39 | Left_A44v | ventral area 44 | DMN |
| 40 | Right_A44v | ventral area 44 | FPN |
| 41 | Left_A14m | medial area 14 | DMN |
| 42 | Right_A14m | medial area 14 | DMN |
| 43 | Left_A12/47o | orbital area 12/47 | DMN |
| 44 | Right_A12/47o | orbital area 12/47 | DMN |
| 45 | Left_A11l | lateral area 11 | LMN |
| 46 | Right_A11l | lateral area 11 | LMN |
| ~~47~~ | ~~Left_A11m~~ | ~~medial area 11~~ | ~~LMN~~ |
| ~~48~~ | ~~Right_A11m~~ | ~~medial area 11~~ | ~~LMN~~ |
| 49 | Left_A13 | area 13 | LMN |
| 50 | Right_A13 | area 13 | LMN |
| 51 | Left_A12/47l | lateral area 12/47 | DMN |
| 52 | Right_A12/47l | lateral area 12/47 | DMN |
| 53 | Left_A4hf | area 4(head and face region) | SMN |
| 54 | Right_A4hf | area 4(head and face region) | SMN |
| 55 | Left_A6cdl | caudal dorsolateral area 6 | DAN |
| 56 | Right_A6cdl | caudal dorsolateral area 6 | DAN |
| 57 | Left_A4ul | area 4(upper limb region) | SMN |
| 58 | Right_A4ul | area 4(upper limb region) | SMN |
| 59 | Left_A4t | area 4(trunk region) | SMN |
| 60 | Right_A4t | area 4(trunk region) | SMN |
| 61 | Left_A4tl | area 4(tongue and larynx region) | VAN |
| 62 | Right_A4tl | area 4(tongue and larynx region) | VAN |
| 63 | Left_A6cvl | caudal ventrolateral area 6 | DAN |
| 64 | Right_A6cvl | caudal ventrolateral area 6 | DAN |
| 65 | Left_A1/2/3ll | area1/2/3 (lower limb region) | SMN |
| 66 | Right_A1/2/3ll | area1/2/3 (lower limb region) | SMN |
| 67 | Left_A4ll | area 4 | SMN |
| 68 | Right_A4ll | area 4 | SMN |
| ~~69~~ | ~~Left_A38m~~ | ~~medial area 38~~ | ~~LMN~~ |
| ~~70~~ | ~~Right_A38m~~ | ~~medial area 38~~ | ~~LMN~~ |
| 71 | Left_A41/42 | area 41/42 | SMN |
| 72 | Right_A41/42 | area 41/42 | SMN |
| 73 | Left_TE1.0 and TE1.2 | TE1_TE1.2 | SMN |
| 74 | Right_TE1.0 and TE1.2 | TE1_TE1.2 | SMN |
| 75 | Left_A22c | caudal area 22 | SMN |
| 76 | Right_A22c | caudal area 22 | SMN |
| 77 | Left_A38l | lateral area 38 | LMN |
| 78 | Right_A38l | lateral area 38 | LMN |
| 79 | Left_A22r | rostral area 22 | DMN |
| 80 | Right_A22r | rostral area 22 | SMN |
| 81 | Left_A21c | caudal area 21 | DMN |
| 82 | Right_A21c | caudal area 21 | DMN |
| 83 | Left_A21r | rostral area 21 | DMN |
| 84 | Right_A21r | rostral area 21 | DMN |
| 85 | Left_A37dl | dorsolateral area37 | DAN |
| 86 | Right_A37dl | dorsolateral area37 | DAN |
| 87 | Left_aSTS | anterior superior temporal sulcus | DMN |
| 88 | Right_aSTS | anterior superior temporal sulcus | DMN |
| 89 | Left_A20iv | intermediate ventral area 20 | LMN |
| 90 | Right_A20iv | intermediate ventral area 20 | LMN |
| 91 | Left_A37elv | extreme lateroventral area37 | DAN |
| 92 | Right_A37elv | extreme lateroventral area37 | DAN |
| 93 | Left_A20r | rostral area 20 | LMN |
| ~~94~~ | ~~Right_A20r~~ | ~~rostral area 20~~ | ~~LMN~~ |
| 95 | Left_A20il | intermediate lateral area 20 | DMN |
| 96 | Right_A20il | intermediate lateral area 20 | LMN |
| 97 | Left_A37vl | ventrolateral area 37 | DAN |
| 98 | Right_A37vl | ventrolateral area 37 | DAN |
| 99 | Left_A20cl | caudolateral of area 20 | FPN |
| 100 | Right_A20cl | caudolateral of area 20 | FPN |
| ~~101~~ | ~~Left_A20cv~~ | ~~caudoventral of area 20~~ | ~~LMN~~ |
| 102 | Right_A20cv | caudoventral of area 20 | LMN |
| 103 | Left_A20rv | rostroventral area 20 | LMN |
| 104 | Right_A20rv | rostroventral area 20 | LMN |
| 105 | Left_A37mv | medioventral area37 | VN |
| 106 | Right_A37mv | medioventral area37 | VN |
| 107 | Left_A37lv | lateroventral area37 | DAN |
| 108 | Right_A37lv | lateroventral area37 | VN |
| 109 | Left_A35/36r | rostral area 35/36 | LMN |
| 110 | Right_A35/36r | rostral area 35/36 | LMN |
| ~~111~~ | ~~Left_A35/36c~~ | ~~caudal area 35/36~~ | ~~LMN~~ |
| 112 | Right_A35/36c | caudal area 35/36 | LMN |
| ~~113~~ | ~~Left_TL~~ | ~~area TL (lateral PPHC~~ | ~~DMN~~ |
| 114 | Right_TL | area TL (lateral PPHC | VN |
| ~~115~~ | ~~Left_A28/34~~ | ~~area 28/34 (EC~~ | ~~LMN~~ |
| ~~116~~ | ~~Right_A28/34~~ | ~~area 28/34 (EC~~ | ~~LMN~~ |
| ~~117~~ | ~~Left_TI~~ | ~~area TI(temporal agranular insular cortex)~~ | ~~LMN~~ |
| ~~118~~ | ~~Right_TI~~ | ~~area TI(temporal agranular insular cortex)~~ | ~~LMN~~ |
| ~~119~~ | ~~Left_TH~~ | ~~area TH (medial PPHC)~~ | ~~VN~~ |
| 120 | Right_TH | area TH (medial PPHC) | VN |
| 121 | Left_rpSTS | rostroposterior superior temporal sulcus | DMN |
| 122 | Right_rpSTS | rostroposterior superior temporal sulcus | DMN |
| 123 | Left_cpSTS | caudoposterior superior temporal sulcus | DMN |
| 124 | Right_cpSTS | caudoposterior superior temporal sulcus | VAN |
| 125 | Left_A7r | rostral area 7 | DAN |
| 126 | Right_A7r | rostral area 7 | DAN |
| 127 | Left_A7c | caudal area 7 | DAN |
| 128 | Right_A7c | caudal area 7 | DAN |
| 129 | Left_A5l | lateral area 5 | DAN |
| 130 | Right_A5l | lateral area 5 | DAN |
| 131 | Left_A7pc | postcentral area 7 | SMN |
| 132 | Right_A7pc | postcentral area 7 | SMN |
| 133 | Left_A7ip | intraparietal area 7(hIP3) | DAN |
| 134 | Right_A7ip | intraparietal area 7(hIP3) | DAN |
| 135 | Left_A39c | caudal area 39(PGp) | VN |
| 136 | Right_A39c | caudal area 39(PGp) | DAN |
| 137 | Left_A39rd | rostrodorsal area 39(Hip3) | FPN |
| 138 | Right_A39rd | rostrodorsal area 39(Hip3) | FPN |
| 139 | Left_A40rd | rostrodorsal area 40(PFt) | DAN |
| 140 | Right_A40rd | rostrodorsal area 40(PFt) | DAN |
| 141 | Left_A40c | caudal area 40(PFm) | DMN |
| 142 | Right_A40c | caudal area 40(PFm) | FPN |
| 143 | Left_A39rv | rostroventral area 39(PGa) | DMN |
| 144 | Right_A39rv | rostroventral area 39(PGa) | DMN |
| 145 | Left_A40rv | rostroventral area 40(PFop) | VAN |
| 146 | Right_A40rv | rostroventral area 40(PFop) | VAN |
| 147 | Left_A7m | medial area 7(PEp) | FPN |
| 148 | Right_A7m | medial area 7(PEp) | DAN |
| 149 | Left_A5m | medial area 5(PEm) | VAN |
| 150 | Right_A5m | medial area 5(PEm) | DAN |
| 151 | Left_dmPOS | dorsomedial parietooccipital sulcus(PEr) | DMN |
| 152 | Right_dmPOS | dorsomedial parietooccipital sulcus(PEr) | VN |
| 153 | Left_A31 | area 31 (Lc1) | DMN |
| 154 | Right_A31 | area 31 (Lc1) | DMN |
| 155 | Left_A1/2/3ulhf | area 1/2/3(upper limb | SMN |
| 156 | Right_A1/2/3ulhf | area 1/2/3(upper limb | SMN |
| 157 | Left_A1/2/3tonIa | area 1/2/3(tongue and larynx region) | SMN |
| 158 | Right_A1/2/3tonIa | area 1/2/3(tongue and larynx region) | SMN |
| 159 | Left_A2 | area 2 | SMN |
| 160 | Right_A2 | area 2 | SMN |
| 161 | Left_A1/2/3tru | area1/2/3(trunk region) | SMN |
| 162 | Right_A1/2/3tru | area1/2/3(trunk region) | SMN |
| 163 | Left_G | hypergranular insula | SMN |
| 164 | Right_G | hypergranular insula | SMN |
| 165 | Left_vIa | ventral agranular insula | DMN |
| 166 | Right_vIa | ventral agranular insula | VAN |
| 167 | Left_dIa | dorsal agranular insula | VAN |
| 168 | Right_dIa | dorsal agranular insula | VAN |
| 169 | Left_vId/vIg | ventral dysgranular and granular insula | VAN |
| 170 | Right_vId/vIg | ventral dysgranular and granular insula | VAN |
| 171 | Left_dIg | dorsal granular insula | SMN |
| 172 | Right_dIg | dorsal granular insula | SMN |
| 173 | Left_dId | dorsal dysgranular insula | VAN |
| 174 | Right_dId | dorsal dysgranular insula | VAN |
| 175 | Left_A23d | dorsal area 23 | DMN |
| 176 | Right_A23d | dorsal area 23 | DMN |
| 177 | Left_A24rv | rostroventral area 24 | FPN |
| 178 | Right_A24rv | rostroventral area 24 | DMN |
| 179 | Left_A32p | pregenual area 32 | DMN |
| 180 | Right_A32p | pregenual area 32 | VAN |
| 181 | Left_A23v | ventral area 23 | DMN |
| 182 | Right_A23v | ventral area 23 | DMN |
| 183 | Left_A24cd | caudodorsal area 24 | VAN |
| 184 | Right_A24cd | caudodorsal area 24 | VAN |
| 185 | Left_A23c | caudal area 23 | VAN |
| 186 | Right_A23c | caudal area 23 | VAN |
| 187 | Left_A32sg | subgenual area 32 | DMN |
| 188 | Right_A32sg | subgenual area 32 | DMN |
| 189 | Left_cLinG | caudal lingual gyrus | VN |
| 190 | Right_cLinG | caudal lingual gyrus | VN |
| 191 | Left_rCunG | rostral cuneus gyrus | VN |
| 192 | Right_rCunG | rostral cuneus gyrus | VN |
| 193 | Left_cCunG | caudal cuneus gyrus | VN |
| 194 | Right_cCunG | caudal cuneus gyrus | VN |
| 195 | Left_rLinG | rostral lingual gyrus | VN |
| 196 | Right_rLinG | rostral lingual gyrus | VN |
| 197 | Left_vmPOS | ventromedial parietooccipital sulcus | VN |
| 198 | Right_vmPOS | ventromedial parietooccipital sulcus | VN |
| 199 | Left_mOccG | middle occipital gyrus | VN |
| 200 | Right_mOccG | middle occipital gyrus | VN |
| 201 | Left_V5/MT+ | area V5/MT+ | VN |
| 202 | Right_V5/MT+ | area V5/MT+ | VN |
| 203 | Left_OPC | occipital polar cortex | VN |
| 204 | Right_OPC | occipital polar cortex | VN |
| 205 | Left_iOccG | inferior occipital gyrus | VN |
| 206 | Right_iOccG | inferior occipital gyrus | VN |
| 207 | Left_msOccG | medial superior occipital gyrus | VN |
| 208 | Right_msOccG | medial superior occipital gyrus | VN |
| 209 | Left_lsOccG | lateral superior occipital gyrus | VN |
| 210 | Right_lsOccG | lateral superior occipital gyrus | VN |
